# DynamicsPLM: A Dynamics-Enriched Small Protein Language Model for Protein Variant Effect Prediction

**DOI:** 10.64898/2026.11.11.765945

**Authors:** Tuan Minh Dao, Amarda Shehu, Anowarul Kabir

## Abstract

Predicting how mutations impact protein function and stability remains a key challenge in computational biology. Current state-of-the-art protein language models (PLMs) primarily scale parameter size on sequence corpora to capture evolutionary constraints but remain blind to the explicit physical dynamics of folded proteins. Conversely, models trained strictly on physical dynamics neglect critical evolutionary sequence statistics. In this paper, we bridge this gap by introducing DynamicsPLM, a highly parameter-efficient 35-million-parameter model that unifies evolutionary and biophysical representation learning. Utilizing a second-stage, multi-task continual pretraining strategy on over 60,000 proteins, we enrich a foundational sequence encoder with explicit residue-level and pairwise molecular dynamics heads while preserving its original evolutionary knowledge. At inference, DynamicsPLM computes zero-shot mutation effects from sequence alone by balancing an evolutionary plausibility score against a structural dynamics disruption score. Across the massive ProteinGym benchmark and natural variants of the Tsuboyama folding stability dataset, DynamicsPLM achieves state-of-the-art accuracy among all models under 100M parameters, and matches or outperforms several billion-scale sequence-based models of its size. Our findings demonstrate that explicit structural-dynamics supervision provides a parameter-efficient alternative to raw scale.

**Availability:** All codes including data preparation scripts, training and validation recipes, and experimental configurations for LoMuS are available at: https://github.com/kabir-ai2bio-lab/DynamicsPLM.

## 1 INTRODUCTION

What a protein molecule does in the living cell, whether it catalyzes a reaction, binds a molecular partner, or provides structural support for the cell, is governed by the three-dimensional structure into which it folds, which in turn is encoded by a protein’s linear sequence of amino acids. A single mutation, the substitution, insertion, or deletion of one amino acid, can leave a protein’s function largely intact or disrupt its folded stability, its folding kinetics, and its biochemical activity. Predicting how a local change in sequence affects a protein’s behavior is the core problem of variant effect prediction, with applications in interpreting disease-associated genetic variants Bromberg et al. (2024), enzyme design Biswas et al. (2021), and rational protein engineering Chubb et al. (2026). Deep mutational scanning (DMS) experiments Fowler and Fields (2014) can measure these functional landscapes directly for thousands to millions of variants in a single assay. However, these experiments are slow, expensive, and specific to a single assay or protein. As a result, the scale of unexplored sequence space makes generalizable, zero-shot prediction across diverse and unseen protein families essential.

To address this challenge, protein language models (PLMs) pretrained with self-supervised masked language modeling (MLM) Devlin et al. (2019) on large natural corpora have emerged as a dominant paradigm Lin et al. (2023); Meier et al. (2021); Notin et al. (2022); Marquet et al. (2022); Cheng et al. (2023); Elnaggar et al. (2022). These architectures implicitly learn the evolutionary constraints encoded in sequence co-occurrence statistics and predict the mutational landscape of natural proteins from sequence alone Meier et al. (2021); Rives et al. (2021); Lin et al. (2023).

However, this sequence-only approach faces two key limitations. First, accuracy gains are primarily due to parameter scaling to hundreds of millions or billions of parameters Lin et al. (2023). Second, sequence-only models are blind to the physical dynamics of the folded structure, which leads them to misjudge mutations that are evolutionarily plausible but structurally destabilizing Bridgham et al. (2009); Starr and Thornton (2016), and variants that are evolutionarily rare yet structurally tolerated Bloom et al. (2005). One line of work adds structural information through multi-modal architectures Kabir and Shehu (2022); Cheng et al. (2024); Li et al. (2024); Su et al. (2024); Tan et al. (2025), but these require experimentally or computationally derived structures at inference. A second line trains directly on biophysical properties from molecular dynamics (MD) simulations Hou et al. (2026); Gelman et al. (2025); because these models train on dynamics rather than sequence, they no longer learn the evolutionary constraints that inform natural amino acid variability.

In this paper, we ask whether a single, small pretrained PLM can be enriched with biophysical dynamics while preserving its evolutionary knowledge, operating from sequence alone at inference without structural inputs or multiple sequence alignments. We introduce **DynamicsPLM**, which subjects a compact ESM2-35M encoder Lin et al. (2023) to a second-stage multi-task continual pretraining phase. By optimizing a joint objective, the model preserves its evolutionary constraints through the original MLM head while internalizing physical constraints through new residue-level and pairwise molecular dynamics prediction heads. At inference, DynamicsPLM scores a mutation by weighing an evolutionary plausibility score against a dynamics disruption score derived from the same enriched representations. Across the ProteinGym deep mutational scanning benchmark Notin et al. (2023) and natural variants of the Tsuboyama folding stability dataset Tsuboyama et al. (2023), DynamicsPLM achieves the best performance among all models under 100M parameters and, on ProteinGym, matches or exceeds sequence-based models up to 86 times its size Lin et al. (2023). These results show that explicit, lightweight dynamics supervision is a parameter-efficient alternative to scaling model size.

## 2 RELATED WORK

### Sequence-Only PLMs

The dominant approach in variant effect prediction relies on scaling unsupervised language modeling over massive natural sequence databases Devlin et al. (2019); Lin et al. (2023); Meier et al. (2021). SOTA models like ESM-2 Lin et al. (2023), Tranception Notin et al. (2022), and ProtTrans Elnaggar et al. (2022) optimize masked or autoregressive objectives to internalize fitness landscapes shaped by natural selection. While highly effective at zero-shot mutation scoring via masked-marginal log-likelihoods Meier et al. (2021), these architectures rely on massive parameter footprints (into the billions) to implicitly capture structural constraints Lin et al. (2023). DynamicsPLM directly contrasts with this paradigm by demonstrating that explicit supervision with biophysical ground truths allows compact representations to bypass the necessity of brute-force sequence scaling.

### Structure-Aware and Multi-Modal PLMs

To address the structural blindness of sequence-only models, multi-modal architectures integrate explicit 3D geometric coordinates into pretraining or fine-tuning pipelines Kabir and Shehu (2022); Cheng et al. (2024); Li et al. (2024); Su et al. (2024); Tan et al. (2025). Models like SaProt Su et al. (2024) and ProSST Li et al. (2024) condition residue embeddings on native backbone structures derived from experimental coordinates or AlphaFold predictions. However, the availability of high-resolution structural inputs at inference introduces a severe practical bottleneck. DynamicsPLM circumvents this dependency by internalizing structural properties directly within its parameter space during pretraining, executing zero-shot inference strictly from sequence inputs.

### Dynamics-Informed Representation Learning

Most closely related to our approach are recent methods that extract biophysical descriptors from molecular dynamics (MD) simulations to train property-aware models Gelman et al. (2025); Hou et al. (2026). ESMDance Hou et al. (2026) established that supervising encoders with residue-level and pairwise MD fluctuations yields embeddings highly sensitive to thermodynamic shifts. These models are optimized strictly on physical dynamics objectives and fail to retain the rich evolutionary sequence statistics captured by PLMs. DynamicsPLM addresses this gap. By utilizing a multi-task, second-stage continual pretraining framework, our model explicitly merges the evolutionary reasoning of MLM with the physical constraints of MD simulations, delivering a parameter-efficient architecture that preserves the strengths of both modalities.

## 3 DATASETS

### 3.1 Second Stage Pretraining & Evaluation Data

We adopt a molecular dynamics (MD) simulation dataset comprising a total of 64, 403 proteins Hou et al. (2026). The dataset is organized into two resolution tiers. High-resolution data consists of all-atom MD simulation trajectories from four sources: mdCATH Mirarchi et al. (2024) (5,392 CATH domains, ∼464 ns × 5 replicates), ATLAS Vander Meersche et al. (2023) (1,516 ordered PDB structures, 100 ns × 3 replicates), GPCRmd Rodríguez-Espigares et al. (2020) (509 membrane proteins, ∼500 ns × 3 replicates), and PED Ghafouri et al. (2024) (382 disordered regions, experimental ensembles). On the other hand, low-resolution data are coarse-grained MD trajectories of human intrinsically disordered regions from IDRome Tesei et al. (2024) (28,058 proteins) and NMA-derived features for ordered PDB structures from Proteinflow Mi et al. (2024) (28,546 proteins). From each protein, we extract 50 residue-level properties per residue and 13 residue-residue biophysical constraint features.

Prior to splitting, we remove proteins whose sequences appear in more than one data source, reducing the 64,403 raw entries to 61,959 unique proteins. To make the train-validation split for the second-order pretraining, we partition the available proteins based on standard sequence identity threshold. Sequences are clustered at 50% identity and 95% of clusters are assigned to train-set, with the remaining 5% held out for evaluation. This yields 58,832 training proteins and 3,127 evaluation proteins, reported in SI Table 2.

**Table 1.**
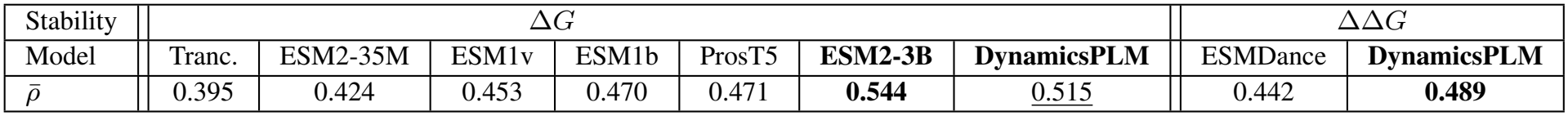
Stability performance on the natural Tsuboyama proteins Tsuboyama et al. (2023) against sequence-based baselines (Spearman *ρ*). DynamicsPLM ranks second overall, behind only ESM2-3B, which has roughly 86× more parameters.

| Stability | ΔG |  |  |  |  |  |  | ΔΔG |  |
|---|---|---|---|---|---|---|---|---|---|
| Model | Tranc. | ESM2-35M | ESM1v | ESM1b | ProsT5 | **ESM2-3B** | **DynamicsPLM** | ESMDance | **DynamicsPLM** |
| ρ̄ | 0.395 | 0.424 | 0.453 | 0.470 | 0.471 | **0.544** | <u>0.515</u> | 0.442 | **0.489** |

### 3.2 Benchmarking Datasets

We leverage two large-scale benchmark datasets to demonstrate the efficacy of our proposed model, DynamicsPLM.

**ProteinGym DMS dataset Notin et al. (2023)** comprises deep mutational scanning (DMS) experiments across a large and diverse set of 217 proteins, measuring the fitness landscapes of amino acid substitutions under various functional assays. We exclude proteins exceeding 1, 022 residues due to the backbone encoder limitation. Additionally, variants having different sequence length from the wild-type due to insertions and/or deletions are excluded. After preprocessing, the benchmark covers 201 proteins having a total of 2, 413, 913 substitution variants. This set can further be studied by the organismal taxons and functional selection types. By organism taxon, there are 87 Human proteins (477, 509 variants), 39 Eukaryote proteins (943, 363 variants), 49 Prokaryote proteins (802, 159 variants), and 26 Virus proteins (190, 882 variants). By functional selection type, the benchmark includes 66 Stability proteins (134, 794 variants), 38 Activity proteins (437, 403 variants), 12 Binding proteins (747, 261 variants), 16 Expression proteins (93, 745 variants), and 69 OrganismalFitness proteins (1, 000, 710 variants). Therefore, ProteinGym benchmarks the broader effectiveness of the proposed second-order embedding enrichment process for mutation effect prediction. Performance is reported as Spearman’s rank correlation (*ρ*) between predicted and experimental scores.

**Tsuboyama dataset Tsuboyama et al. (2023)** is a mega-scale protein folding stability benchmark containing experimentally measured Δ*G* and ΔΔ*G* values for single- and multi-site substitutions across 479 proteins, encompassing both natural and *de novo* designed sequences. We use the fitness label (`dG_ML` and `ddG_ML`) as the ground-truth folding stability value. We exclude any rows with missing `ddG_ML` values (no missing for `dG_ML`) and all insertion/deletion, retaining only substitution mutations. After preprocessing, the benchmark comprises 412 proteins of 527, 794 substitution variants in total for ΔΔ*G* evaluation. Proteins are further classified into 266 natural proteins (404, 685 variants) and 146 *de novo* designed proteins (123, 109 variants). Performance is measured by Spearman’s rank correlation *ρ* between predicted scores and experimental ΔΔ*G* values, reported per protein and averaged across all proteins.

## 4 METHODOLOGY

### 4.1 Model Architecture

We aim to enrich the representations of a small pretrained PLM to enhance embedding quality for challenging downstream prediction tasks. To achieve this, we enforce the backbone model to learn structural and biophysical information explicitly while preserving its previously learned evolutionary information. Specifically, we introduce a second-order continual pretraining strategy that optimizes the model using a multi-task learning objective. The model architecture is illustrated in Fig. 1.

**Figure 1.**
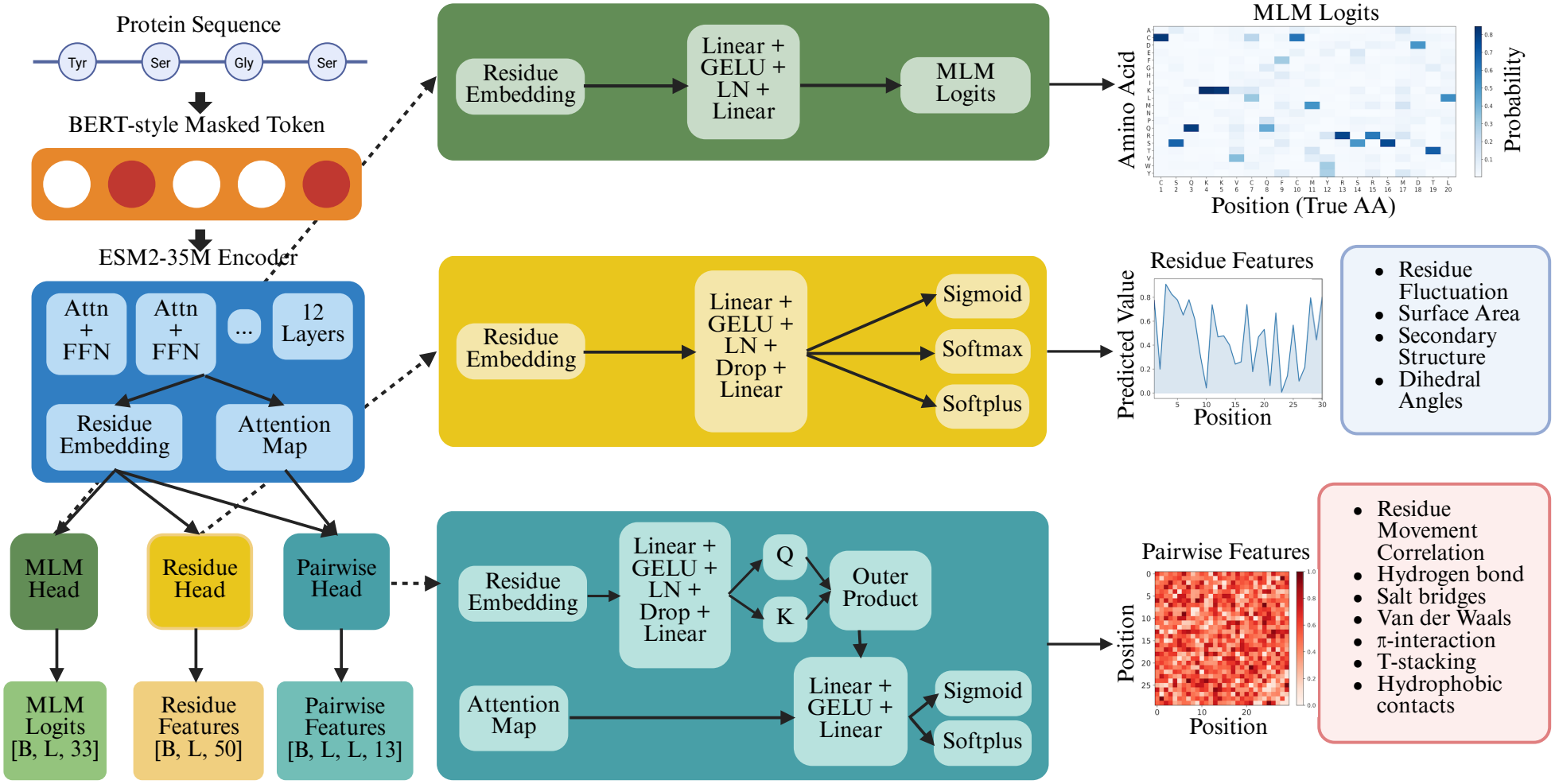
Overview of DynamicsPLM. A protein sequence is tokenized and masked following the BERT Devlin et al. (2019) strategy before being processed by the ESM-2 Lin et al. (2023) 35M backbone encoder. Per-residue embeddings and multi-layer attention maps are passed to three jointly trained output heads: an MLM head predicting masked amino acid probabilities, a residue dynamics head predicting residue-level features (RMSF, SASA, secondary structure, and torsion angles), and a pairwise dynamics head predicting pairwise interaction features (van der Waals contacts, hydrogen bonds, salt bridges, hydrophobic contacts, and residue movement correlations). Example outputs of each head are shown on the right.

We leverage a small pretrained transformer-based PLM, ESM-2 35M Lin et al. (2023) variant, as our backbone encoder architecture. DynamicsPLM enriches the PLM with molecular dynamics information through multi-task supervision, then leverages the enriched representation for zero-shot mutation effect prediction. By design, the model takes only protein sequences as input. The sequence is tokenized using the ESM-2 tokenizer and a random 15% of residue tokens are masked following the BERT Devlin et al. (2019) strategy. The encoder processes the masked sequence and produces per-residue contextual embeddings alongside multi-layer attention maps. These representations are passed to three output heads: an MLM head, a residue-level dynamics head, and a pairwise dynamics head. The entire parameter space is then optimized through a joint loss function encompassing all three tasks using standard backpropagation. Implementation details, hyperparameters and other settings are detailed in the Supplementary Information (SI) 5.

– *MLM head* predicts the probability (or logits) of an amino acid being at the *i*-th position given the input sequence context to the model. It reflects the sequence-level evolutionary constraints.
– *Residue head* deploys a multi-layer perceptron (MLP) after the final backbone encoder layer and predicts residue-level dynamic properties of an input protein molecule. Following Hou et al. (2026), this head predicts 50 residue-level features (listed in SI Table 3).
*Pairwise head* captures the residue-reside biophysical constraints, such as van der Waals interactions, hydrogen bonds, hydrophobic contacts, salt bridges, pairwise movement correlations, and NMA-derived pairwise features (13 total following Hou et al. (2026), in SI Table 3). Residue embeddings are transformed into query and key representations and combined via outer-product operations to form pairwise embeddings, which are concatenated with attention maps from all encoder layers. A linear projection then predicts the pairwise dynamic properties.

Example outputs of each head are depicted in Fig. 1. This ensures that the encoder will internalize structural and biophysical information while preserving or improving the learned evolutionary sequence knowledge.

### 4.2 Objective Function

DynamicsPLM parameter space is optimized with a multi-task learning objective composing sequence- and dynamics-learning loss functions, defined in Eq. 1.

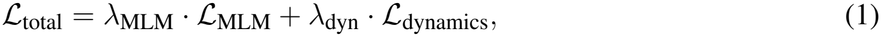

where *λ*_MLM_ and *λ*_dyn_ weight the relevance between evolutionary constraints and molecular dynamics, respectively. ℒ_MLM_ is the standard MLM cross-entropy loss over the masked tokens. For a sequence with ℳ masked positions, ℒ_MLM_ is defined as:

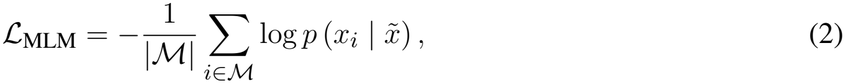

where *x*_*i*_ is the original amino acid token and 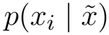 is the softmax probability assigned to it by the MLM head given the masked input 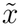.

The dynamics loss ℒ_dynamics_ follows Hou et al. Hou et al. (2026): for each feature *f* , a per-protein RMSE is computed over valid (non-sentinel) residue positions and residue pairs, and the per-feature losses are combined through a source-specific weight table *w*_*f,s*_ that balances contributions across features and data sources (full formulation in SI 5.4).

We studied three variants of DynamicsPLM under different (*λ*_MLM_, *λ*_dyn_) weightings in Eq. 1: 0.3/0.7, 0.5/0.5, and 0.7/0.3, and chose the best 0.7/0.3 configurations in all zero-shot mutation effect prediction (details in Section 6.3, SI Table 4 and SI Table 5).

### 4.3 Inference: Zero-shot Mutation Effect Prediction

We design a zero-shot mutation inference considering both sequence-level evolutionary plausibility from the MLM head and structural disruption from the dynamics heads, without any task-specific fine-tuning.

– *MLM scoring*. We adopt the masked-marginal protocol Rives et al. (2021) to compute the mutation effect from the MLM head. For a sequence of length *L*, we perform *L* forward passes, masking one position at a time, and record the log-probability of each token under the MLM head. The MLM score for a variant *v* relative to the wild-type *w* is:

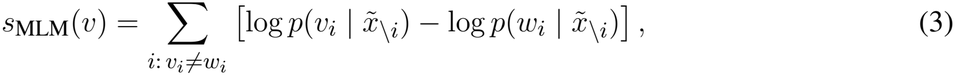

where 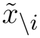 denotes the sequence context with *i*-th position masked. This formulation can handle both single- and multi-site substitutions naturally.
– *Dynamics disruption scoring*. One forward pass is run on both the wild-type and mutant sequences to obtain predictions from the residue and pairwise heads. The per-feature relative disruption is computed as averaged over residue positions or residue pairs:

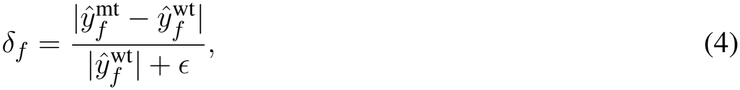

where *ϵ* = 10^−2^ guards against division by near-zero values. Per-feature scores are then quantile-normalised across all variants to equalise feature scales and aggregated via the geometric mean into a single disruption score *s*_dyn_(*v*).
– *Combined scoring*. A higher MLM score indicates evolutionary plausibility whereas a higher dynamics disruption score indicates structural destabilisation. Therefore, we combine these two signals by subtracting dynamics disruptions from the evolutionary plausibilities as:

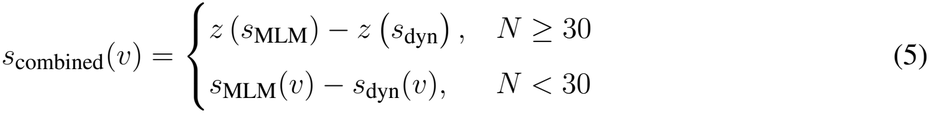

where *z*(·) denotes z-score normalization across all variants of a given protein and *N* is the number of mutated variants available for that protein. Z-score normalization requires a sufficiently large sample to yield stable mean and variance estimates; for proteins with fewer than 30 variants we instead subtract the raw scores directly.

## 5 IMPLEMENTATION DETAILS

### 5.1 Data Processing

We extract the raw amino acid sequence from the training dataset, which serves as the main input to our model. The sequence is tokenized using the ESM-2 tokenizer with CLS and EOS tokens appended. We apply BERT-style masking at *p* = 0.15, where 80% of selected positions are replaced with [MASK], 10% with a randomly sampled amino acid token, and 10% are left unchanged. Special tokens (CLS, EOS, PAD) are excluded from masking eligibility. For sequences exceeding the training maximum length, we draw a random window during training and use a fixed N-terminal window during validation for reproducibility. Variable-length batches are padded to the batch maximum, with residue and pairwise features padded using the −1 sentinel. We employ three independent DistributedSampler-backed DataLoaders that cycle continuously over the three source groups (ATLAS/GPCRmd/PED/mdCATH, IDRome, Proteinflow), each contributing one batch per gradient step. DataLoader random seeds are set to `n_update + device_id` for distributed reproducibility.

### 5.2 Model Training and Evaluation

We train the model using PyTorch DDP via `torchrun` with the NCCL backend and FP16 automatic mixed precision with gradient scaling. The optimizer is AdamW (*β*_1_ = 0.9, *β*_2_ = 0.98, *ε* = 10^−8^, weight decay = 0.01). The encoder and MLM head use a peak learning rate of 10^−5^; we apply a 10× multiplier (peak 10^−4^) to the dynamics head parameters to compensate for training from random initialization. The learning rate schedule consists of a linear warmup over 700 steps, followed by linear decay to 10% of peak over 90% of training, then a fixed floor. We freeze the encoder for the first 2 epochs so the dynamics heads reach a reasonable state before encoder updates begin. Training proceeds in two stages: sequences of length ≤ 256 for the first 15,000 updates, then ≤ 1,024 for the remaining 5,000 (20,000 total). The effective batch size is 32 throughout (batch 16, gradient accumulation 2 in the short phase; batch 2, accumulation 16 in the long phase). Gradients are value-clipped at 0.5. Loss weights are *λ*_mlm_ = 0.7 and *λ*_dyn_ = 0.3. Within ℒ_dynamics_, per-feature weights *w*_*f,s*_ are precomputed from training proteins following SeqDance Hou et al. (2026). Each feature is weighted by the inverse of its standard deviation across training proteins; pairwise interaction features (van der Waals, hydrogen bonds, hydrophobic contacts, salt bridges, and pi interactions) are cube-root transformed before standard deviation estimation to reduce frequency skew. The total pairwise loss is scaled to three times the total residue loss, and within the pairwise group the nine interaction features are collectively weighted three times more than the correlated-motion feature. Contributions from the three source groups (high-resolution MD, IDRome, and Proteinflow NMA) are equalized across sources. We save checkpoints every 1,000 updates and maintain latest checkpoints for resumption. For evaluation, each periodic checkpoint (`update_1000` through `update_20000`) is scored on the 3,127 held-out test proteins to identify the best checkpoint with the lowest dynamics loss, computed as per-source mean RMSE over all dynamic features under `torch.no_grad()` on a single GPU with N-terminal windowing.

### 5.3 Potential data leakage assessment

We performed an exact sequence-identity check between the DynamicsPLM training set and each benchmark. For the ProteinGym DMS substitution benchmark (201 proteins with sequence length ≤ 1,022 aa), 11 wildtype sequences appear verbatim in the training set: 10 in the training split (B2L11_HUMAN, CCDB_ECOLI, GLPA_HUMAN, P84126_THETH, R1AB_SARS2, SPA_STAAU, SPTN1_CHICK, SRBS1_HUMAN, YAP1_HUMAN, and one additional CCDB_ECOLI entry) and one in the held-out evaluation split (P53_HUMAN). For the Tsuboyama stability benchmark (412 proteins), 6 wildtype sequences overlap with the training split (1LP1, 1TUD, 1YRF, 2JN4, 2O2W, 3I35), all sourced from mdCATH or Proteinflow. These overlapping proteins represent 5.5% and 1.5% of each benchmark, respectively.

### 5.4 Dynamics Loss Formulation

Following Hou et al. Hou et al. (2026), the dynamics loss is the source-weighted sum of per-feature residue and pairwise RMSE terms. For each feature *f* , the residue loss averages the per-protein RMSE over valid positions *V*_*p*_ (sentinel value −1 excluded):

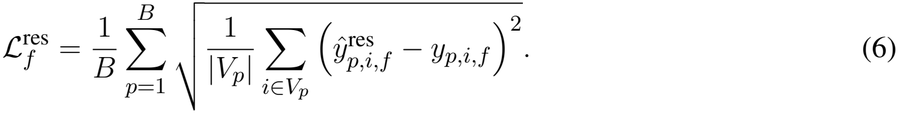

The pairwise loss is defined analogously over valid residue pairs 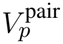:

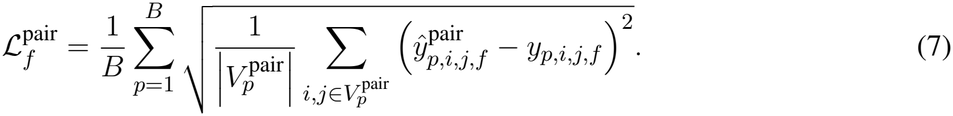

Per-feature losses are combined through a source-specific weight table *w*_*f,s*_:

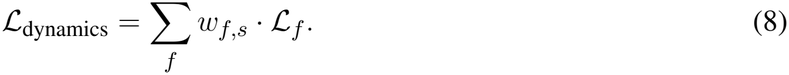

## 6 RESULTS AND DISCUSSION

### 6.1 Benchmarking on the ProteinGym DMS Dataset

We benchmark DynamicsPLM against 46 published models on the ProteinGym Notin et al. (2023) DMS dataset, covering 201 proteins. The performance assessment is summarized in Fig. 2.

**Figure 2.**
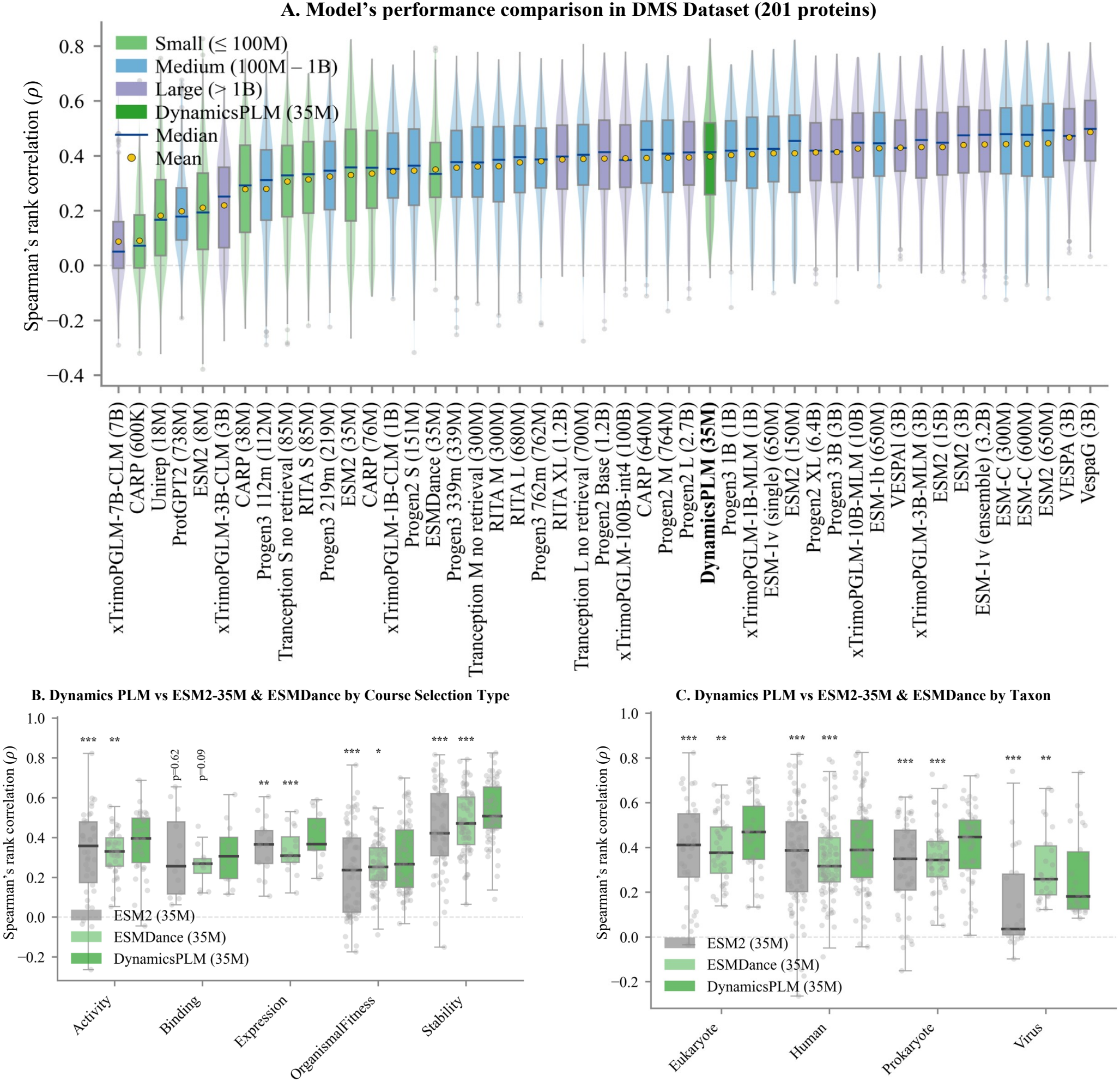
Performance of DynamicsPLM on the ProteinGym DMS substitution benchmark (201 proteins, seq_len ≤ 1022). (A) Comparison of Spearman’s rank correlation (*ρ*) box plots overlaying on violin plots to demonstrate the *ρ* distribution across 201 proteins for 47 sequence-based models, sorted by mean *ρ*. DynamicsPLM (35M) achieves a mean *ρ* of 0.398 and a median *ρ* of 0.414, ranking 1st among all small-scale models (≤ 100M parameters) and 19th overall. (B) Comparison of Spearman’s rank correlation (*ρ*) by per-course selection type for ESM2-35M, ESMDance, and DynamicsPLM. Mean/median *ρ* values achieved by ESM2-35M, ESMDance, and DynamicsPLM are 0.319*/*0.357, 0.324*/*0.329, and 0.375*/*0.395 for Activity; 0.307*/*0.255, 0.260*/*0.268, and 0.317*/*0.306 for Binding; 0.357*/*0.365, 0.335*/*0.308, and 0.398*/*0.367 for Expression; 0.229*/*0.236, 0.262*/*0.252, and 0.299*/*0.266 for OrganismalFitness; and 0.439*/*0.421, 0.479*/*0.471, and 0.528*/*0.507 for Stability, respectively. (C) Similar comparison grouped by organism taxon. Mean/median *ρ* values achieved by ESM2-35M, ESMDance, and DynamicsPLM are 0.389*/*0.411, 0.394*/*0.377, and 0.451*/*0.469 for Eukaryote; 0.364*/*0.386, 0.342*/*0.317, and 0.405*/*0.389 for Human; 0.323*/*0.349, 0.354*/*0.344, and 0.411*/*0.447 for Prokaryote; and 0.141*/*0.035, 0.311*/*0.258, and 0.268*/*0.181 for Virus, respectively. Asterisks in panels B and C denote paired Wilcoxon signed-rank significance against DynamicsPLM (^***^*p <* 0.001, ^**^*p <* 0.01, ^*^*p <* 0.05).

#### Overall performance comparison

DynamicsPLM ranks first among all small models (defined by having ≤100M parameters) and 19th overall out of 47 models, illustrated in Fig. 2A, sorted by the mean Spearman’s rank correlation metric 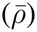. DynamicsPLM achieves a 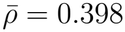 (median 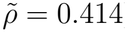) across all 201 proteins. When evaluated against all 47 models, scaling up to 3B parameters, DynamicsPLM outperforms 28 models. Note that six out of these 28 models are at billion scale and 13 fall within the 100M to 1B range. In particular, DynamicsPLM outperforms Tranception L (700M, 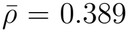), RITA XL (1.2B, 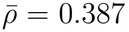), Progen3 (762M, 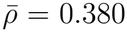), and xTrimoPGLM-1B-CLM (1B, 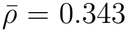). The margin over ESMDance 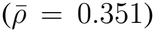 and ESM2-35M 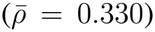, which share the same backbone and similar parameter count, confirms that dynamics supervision provides a consistent benefit over pure masked-language-model pretraining on similar model scale.

#### Per-course selection type analysis

Fig. 2B further breaks down per-protein Spearman’s rank correlations for ESM2-35M, ESMDance, and DynamicsPLM across the five ProteinGym DMS functional assay categories (Activity, Binding, Expression, OrganismalFitness and Stability). DynamicsPLM significantly outperforms both baselines on four of the five categories. The strongest and most significant gain occurs on Stability (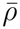: ESM2-35M = 0.439, ESMDance = 0.479, DynamicsPLM = 0.528, *p <* 10^−8^ against both baselines), consistent with the dynamics heads directly modeling the biophysical folding properties that govern thermostability assays. The improvement is also highly significant on OrganismalFitness (*p* = 6.6 × 10^−8^ versus ESM2-35M), despite being the lowest-scoring category overall (ESM2-35M 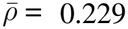). This result suggests that dynamics supervision provides the greatest benefit where the evolutionary signal alone is weak. The one exception is Binding, the smallest category by assay count (*n* = 12), where DynamicsPLM shows no significant advantage over either baseline statistically (*p >* 0.05). With so few assays, the paired test lacks sufficient power to resolve a difference at this scale. However, a side-by-side visual box-plot comparison in Fig. 2B shows an evidence of improvement over both by the DynamicsPLM. This improvement pattern across all assay categories indicates that dynamics supervision contributes most where the assayed phenotype is structurally or biophysically grounded. In these cases, the dynamics-disruption score complements rather than substitutes for the evolutionary signal captured by the MLM head.

#### Per-taxon analysis

DynamicsPLM significantly outperforms ESM2-35M on all four taxa, with p-values of 2.1 × 10^−4^ for Eukaryote, 9.6 × 10^−6^ for Human, 1.8 × 10^−8^ for Prokaryote, and 4.2 × 10^−7^ for Virus (Fig. 2C). It also achieves superior performance over ESMDance on Eukaryote, Human, and Prokaryote proteins with p-values of 1.9 × 10^−3^, 7.0 × 10^−9^, and 3.1 × 10^−4^, respectively. When compared with all 47 models, DynamicsPLM ranks 1st among all models with ≤100M parameters on each of these three taxa and places 11th, 21st, and 17th overall (SI Fig. 6).

The one exception is viral proteins, where ESMDance edges out DynamicsPLM (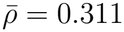 vs. 0.268, *p* = 3.2 × 10^−3^, Fig. 2C), though DynamicsPLM still nearly doubles the ESM2-35M backbone (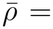 = 0.141 → 0.268). The cause is specific to the combination rather than the dynamics signal: on viral proteins our dynamics-only score already matches ESMDance’s, but the rapid divergence of viral sequences Huot et al. (2026) leaves the MLM component weak, and fusing it with the dynamics score drags the combined correlation below the dynamics-only baseline (per-component breakdown in SI Table 4). The benefit of joint supervision is therefore strongest on eukaryotic, human, and prokaryotic proteins, where evolutionary conservation remains informative.

### 6.2 Benchmarking on Folding Stability

Protein stability is a key determinant for understanding protein function and critical engineering tasks. We analyze our proposed DynamicsPLM in depth across two stability benchmark datasets.

#### 6.2.1 Using ProteinGym DMS Stability benchmark

DynamicsPLM achieves the second-highest mean Spearman’s correlation of all 47 sequence-based models evaluated on the 66 stability assays in ProteinGym 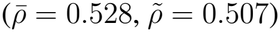, trailing only VespaG, a 3B-parameter model with ∼ 86 times more parameters 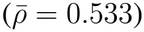 (Fig. 3). DynamicsPLM outperforms several other substantially larger models on this category, including ESM2-3B (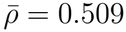, also 86 times larger), ESM2-650M (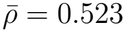, 19 times larger), and ESM-C 300M (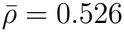, 9 times larger). Among models with ≤100M parameters, DynamicsPLM ranks 1st by a wide margin, exceeding ESMDance 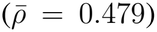 and the ESM2-35M backbone 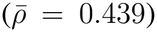 by 0.05 and 0.09 points, respectively. This result indicates that the biophysical properties learned by the dynamics heads, particularly residue burial, backbone rigidity, and pairwise contact energetics, are highly informative for thermostability assays, and that combining them with evolutionary information from the MLM head allows a 35M-parameter sequence-only model to match or exceed the performance of models with substantially larger parameter counts.

**Figure 3.**
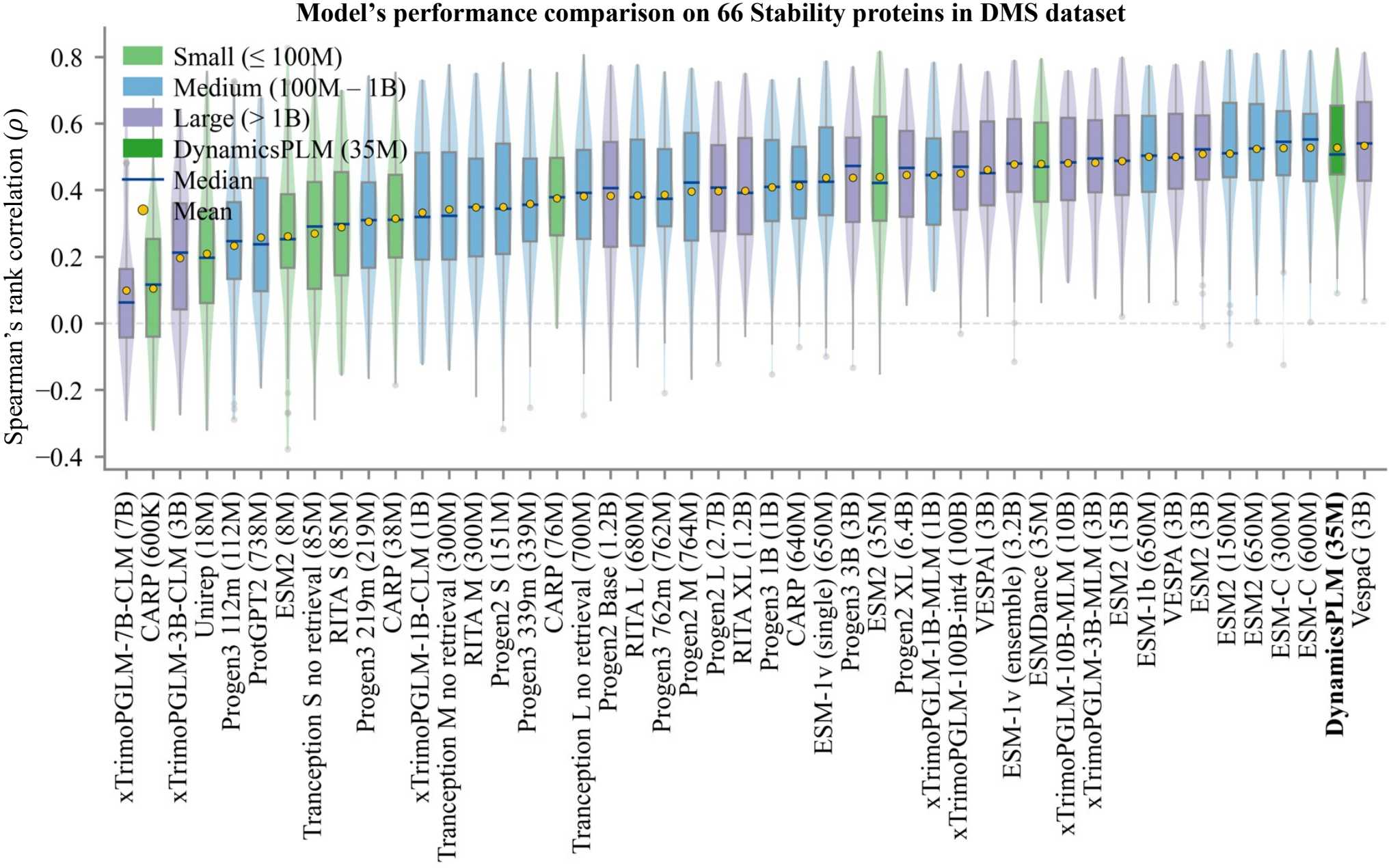
Performance comparison of DynamicsPLM on ProteinGym DMS stability prediction tasks across 47 sequence-based models. Spearman’s rank correlation *ρ* distribution is plotted across 66 proteins. Distribution plots are sorted by mean *ρ*. DynamicsPLM (35M) achieves a mean *ρ* of 0.528 and a median *ρ* of 0.507, ranking 1st among all small-scale models (≤100M parameters) and 2nd overall.

#### 6.2.2 Evaluation on Natural Proteins in Tsuboyama Dataset

DynamicsPLM and ESMDance are now evaluated directly using the same zero-shot scoring protocol, while baseline scores for the remaining sequence-based architectures are sourced from the ProMEP benchmark Cheng et al. (2024). DynamicsPLM consistently outperforms substantially larger sequence-based architectures, trailing only ESM2-3B which possesses ∼ 86 times more parameters Lin et al. (2023), reported in Table 1, in estimating Δ*G* while outperforms ESMDance by a large margin in estimating ΔΔ*G*.

On the 404,685 substitution variants across the 266 natural proteins, DynamicsPLM’s combined score correlates with experimental ΔΔ*G* at *ρ* = 0.489, with low absolute error (MAE = 1.657 and RMSE = 1.987. The per-protein margins in Fig. 4 reveal complementarity rather than one signal dominating: ESMDance beats the ESM2-35M backbone on only 43% of proteins, yet DynamicsPLM beats ESM2-35M on 70% and ESMDance on 76%, because the evolutionary and dynamics heads are informative on different proteins.

**Figure 4.**
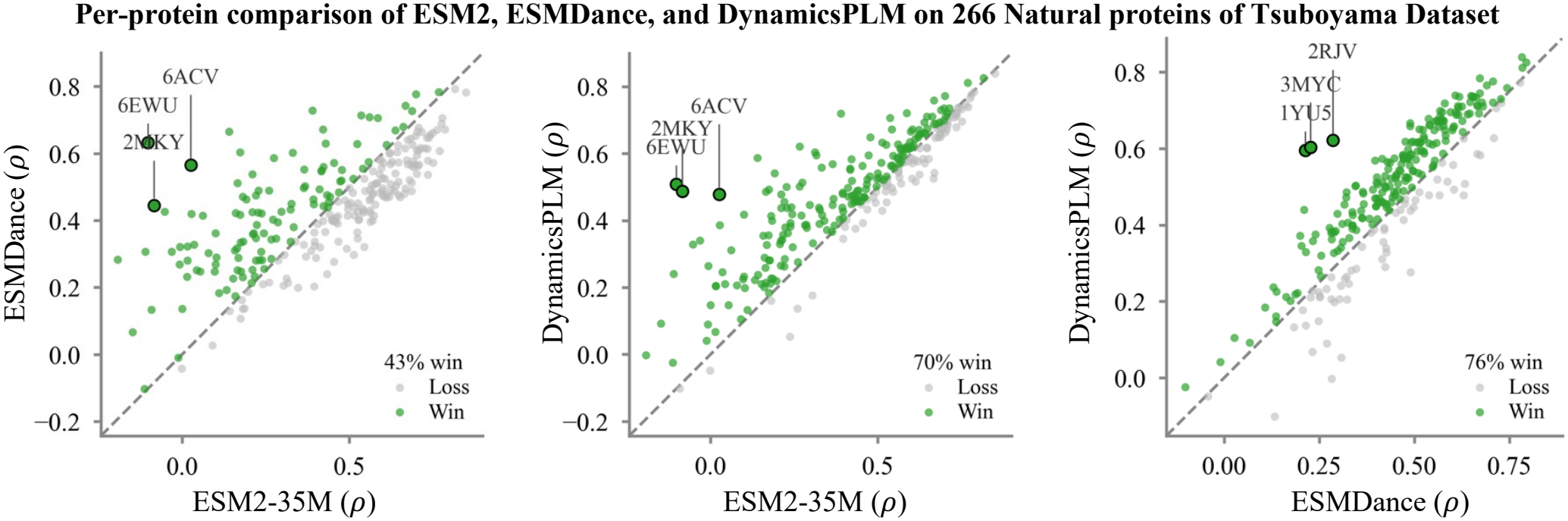
Per-protein Spearman *ρ* on the 266 natural Tsuboyama proteins; points above the diagonal favor the *y*-axis model. ESMDance beats ESM2-35M on 43% of proteins, while DynamicsPLM beats ESM2-35M on 70% and ESMDance on 76%, showing the two heads are informative on different proteins.

#### 6.2.3 Case study: Stability Comparison on Natural Tsuboyama Protein PDB-ID:1YU5

The protein 1YU5 (2, 569 mutations) illustrates the performance superiority of DynamicsPLM over ESM2-35M and ESMDance concretely in Fig. 5. ESM2-35M and ESMDance achieve only weak rank correlation with the true stability ranking (*ρ* = 0.316 and *ρ* = 0.213, respectively), whereas DynamicsPLM nearly doubles this correlation (*ρ* = 0.595). Since neither the evolutionary signal alone nor the dynamics signal alone is sufficient on this protein, the combined score’s advantage here reflects the complementary information captured by the two heads rather than a gain attributable to either signal in isolation.

**Figure 5.**
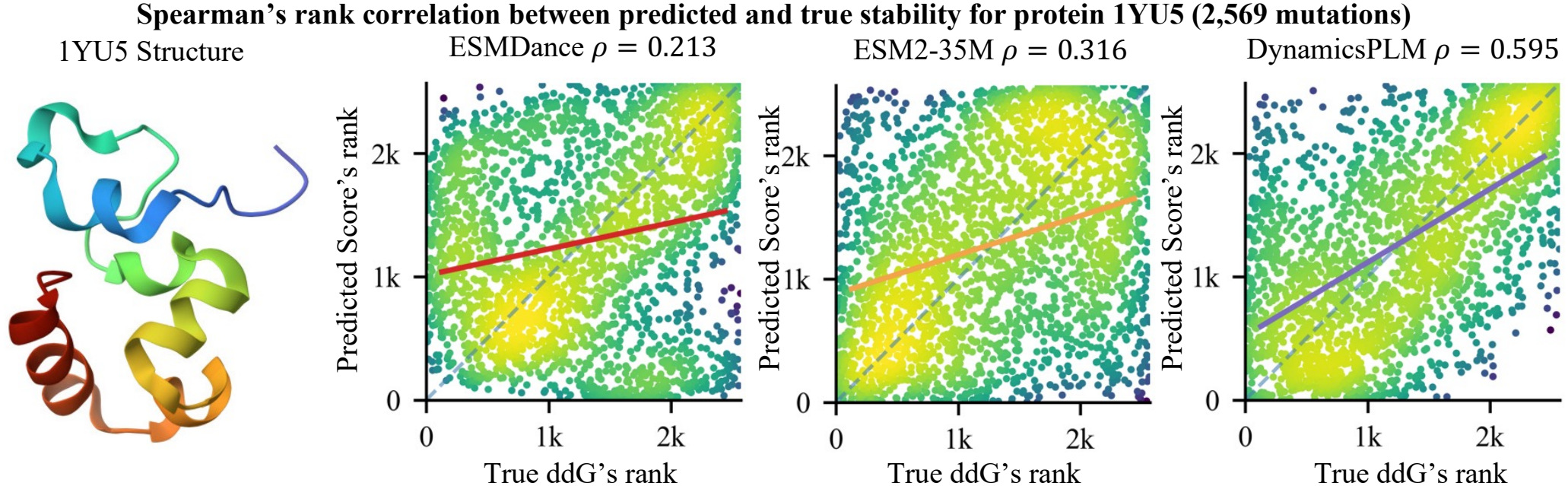
Spearman’s rank correlation between predicted and true stability rank for protein 1YU5 (2,569 mutations): ESMDance *ρ* = 0.213, ESM2-35M *ρ* = 0.316, and DynamicsPLM *ρ* = 0.595.

### 6.3 Ablation on DynamicsPLM configurations

To balance the MLM and dynamics objectives during continual pretraining, we trained three variants of DynamicsPLM under different (*λ*_MLM_, *λ*_dyn_) weightings in Eq. (1): 0.3*/*0.7, 0.5*/*0.5, and 0.7*/*0.3. For each variant, SI Table 4 reports three Spearman correlations against experimental DMS fitness scores. MLM *ρ* is the correlation between the masked-marginal MLM score (Eq. (3)) and fitness. Dyn. *ρ* is the correlation between the dynamics disruption score (Eq. (4)) and fitness. Comb. *ρ* is the correlation between their z-score combination (Eq. (5)) and fitness. Across all 201 proteins, the 0.7 MLM + 0.3 Dyn weighting achieves the highest combined correlation (Comb.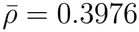, Comb.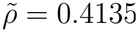), ahead of the 0.5*/*0.5 weighting (0.3953*/*0.4069) and the 0.3*/*0.7 weighting (0.3896*/*0.3960). This configuration also remains the best or near-best within every taxon subgroup. These results indicate that, even though the dynamics heads supply most of the training signal, weighting the MLM objective more heavily during continual pretraining better preserves the evolutionary information needed for zero-shot mutation scoring. Thus, we adopt *λ*_MLM_ = 0.7 and *λ*_dyn_ = 0.3 as the final DynamicsPLM configuration in benchmarking and analysis.

To confirm that this configuration generalizes beyond the ProteinGym ablation, we repeated the loss-weight test on the independent Tsuboyama dataset, using the same MLM *ρ*, Dyn. *ρ*, and Comb. *ρ* metrics defined by Eq. (3), Eq. (4), and Eq. (5). Across all 412 proteins, the 0.7 MLM + 0.3 Dyn weighting again achieves the highest combined correlation (Comb.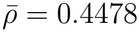, Comb.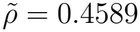, SI Table 5), confirming the configuration selected from the ProteinGym ablation.

On the 266 natural proteins, the three weightings perform within 0.0014 of each other on Comb. 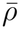 (0.4892 to 0.4906) and within 0.0016 on Comb. 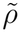 (0.5079 to 0.5095), so the choice of *λ*_MLM_ makes little practical difference on this subset. On the 146 de novo proteins, by contrast, the 0.7 MLM + 0.3 Dyn weighting improves Comb. 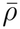 from 0.3538 to 0.3724 relative to the 0.3 MLM + 0.7 Dyn weighting, a much larger gap than on natural proteins. Increasing the MLM weight during training therefore makes little difference on natural proteins, where all three configurations are already strong, but provides a meaningful benefit on de novo proteins, which is what drives the overall preference for *λ*_MLM_ = 0.7 established earlier in this section.

## 7 CONCLUSION

We presented DynamicsPLM, a sequence-only PLM that enriches a compact pretrained encoder with explicit molecular dynamics supervision through a multi-task continual pretraining objective. By coupling MLM with residue-level and pairwise dynamics predictions, our architecture internalizes crucial biophysical properties natively within its latent space. At inference, the model performs zero-shot mutation effect prediction from sequence alone by balancing a masked-marginal evolutionary plausibility score against a structural dynamics disruption score without requiring task-specific fine-tuning. Evaluating our model across the massive ProteinGym dataset and natural variants of the Tsuboyama folding stability benchmark demonstrates that DynamicsPLM consistently ranks first among evaluated models with no more than 100M parameters. Furthermore, it matches or exceeds the accuracy of several purely sequence-based models up to 86 times its size, establishing dynamics-aware continual pretraining as a parameter-efficient alternative to brute-force model scaling.

Even with dynamics supervision, DynamicsPLM’s zero-shot score still leans heavily on evolutionary history, because the fixed weight combination in Eq. (5) always folds in the evolutionary plausibility term regardless of how informative it is. When that history is weak or absent, such as rapidly evolving viral proteins, or *de novo* designed sequences with no natural homologs, the language modeling head adds little signal, and combining it with the dynamics score can lower the correlation below using the dynamics score alone. The clearest next step is to replace the fixed combination with a learned mixing ratio, predicted from sequence-level features such as homolog depth or conservation, so the model can down-weight the evolutionary component when it is uninformative.

## CONFLICT OF INTEREST STATEMENT

No competing interest is declared.

## DATA AVAILABILITY STATEMENT

No new data has been produced during this work. The data used in the model development and validation process can be found in Notin et al. (2023); Tsuboyama et al. (2023). All codes, including data download and preparation scripts, training and validation recipes, and experimental configurations for LoMuS are available at: https://github.com/kabir-ai2bio-lab/DynamicsPLM.

## Supplementary Information

## 8 TRAIN AND EVALUATION DATASET DETAILS

**Table 2.**
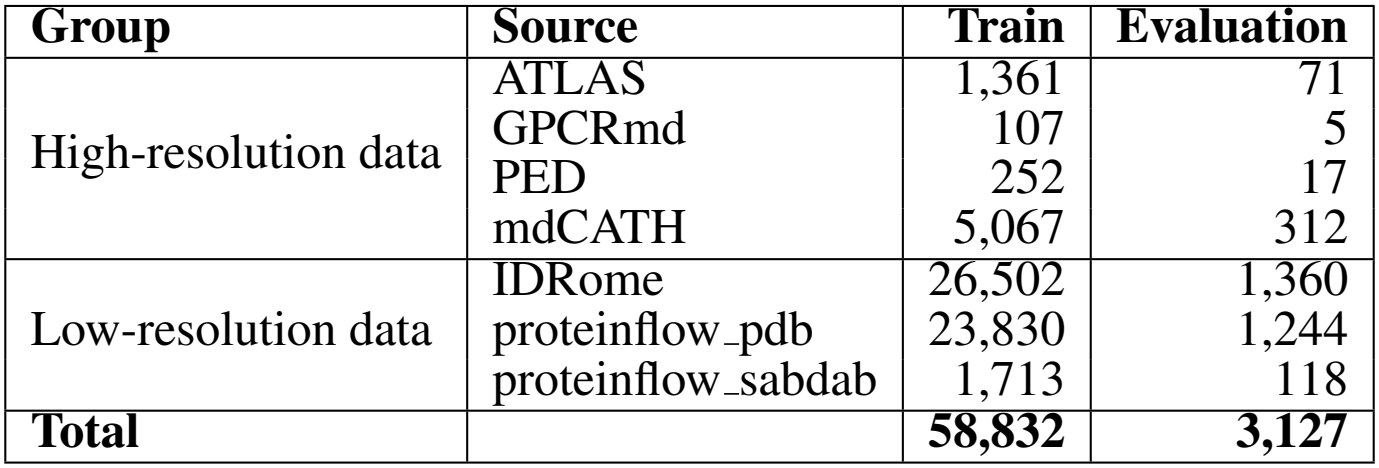
Number of Proteins per Data Source in the Train and Evaluation Sets. We source this data from ESMDance Hou et al. (2026).

## 9 RESIDUE-LEVEL AND PAIRWISE DYNAMIC FATURES

**Table 3.**
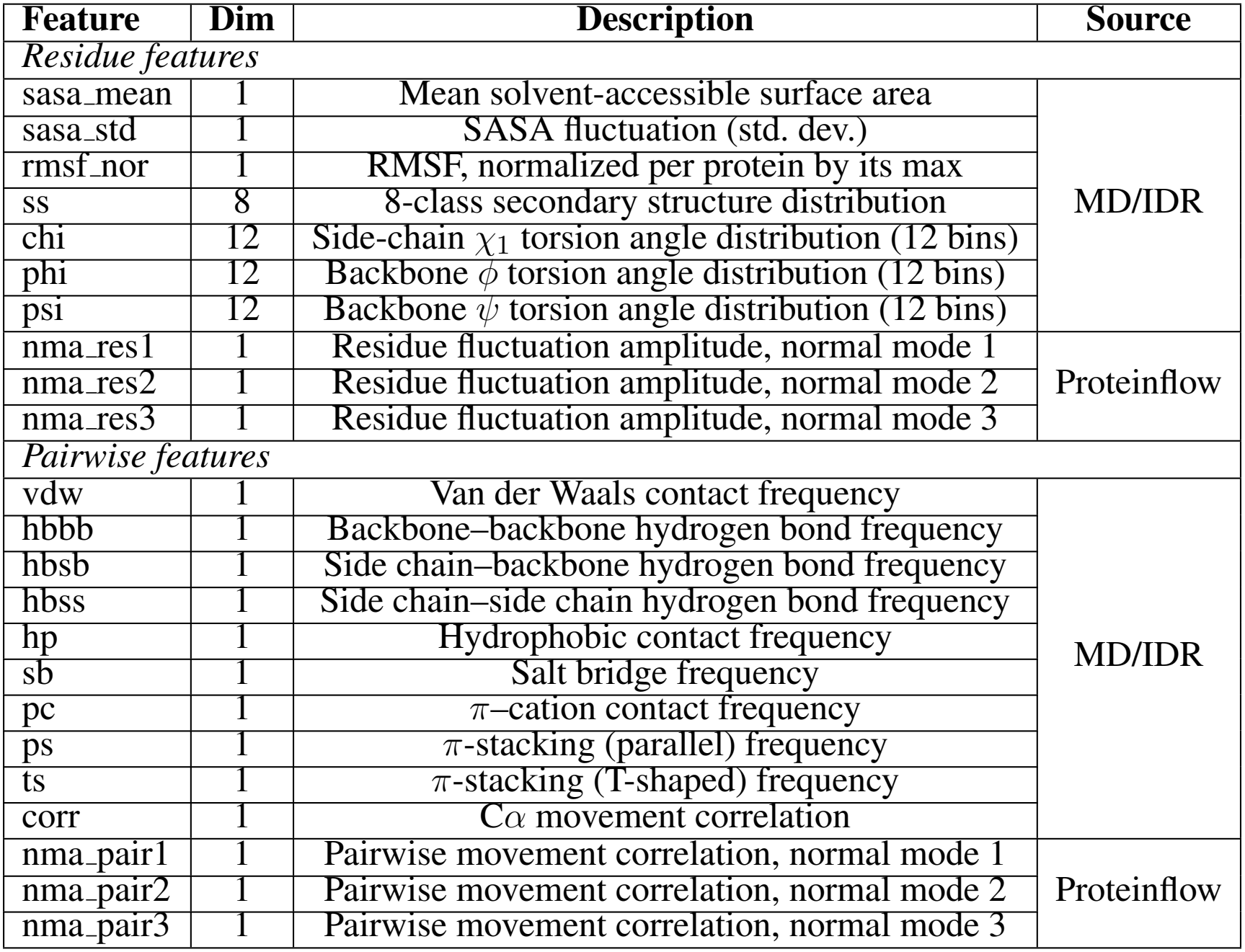
Residue-level (50) and pairwise dynamic (13) features used to enrich DynamicsPLM, with their dimensionality and data source. MD/IDR sources (ATLAS, GPCRmd, PED, mdCATH, IDRome) supply the SASA, RMSF, secondary structure, torsion angle, and pairwise contact features; Proteinflow (pdb and SAbDab structures) supplies normal-mode-analysis-derived features only. For a given protein, features unavailable from its source carry the sentinel value (−1) and are excluded from the loss.

## 10 PER-TAXON MODEL PERFORMANCE COMPARISON ON PROTEINGYM DMS SUBSTITUTION BENCHMARK

**Figure 6.**
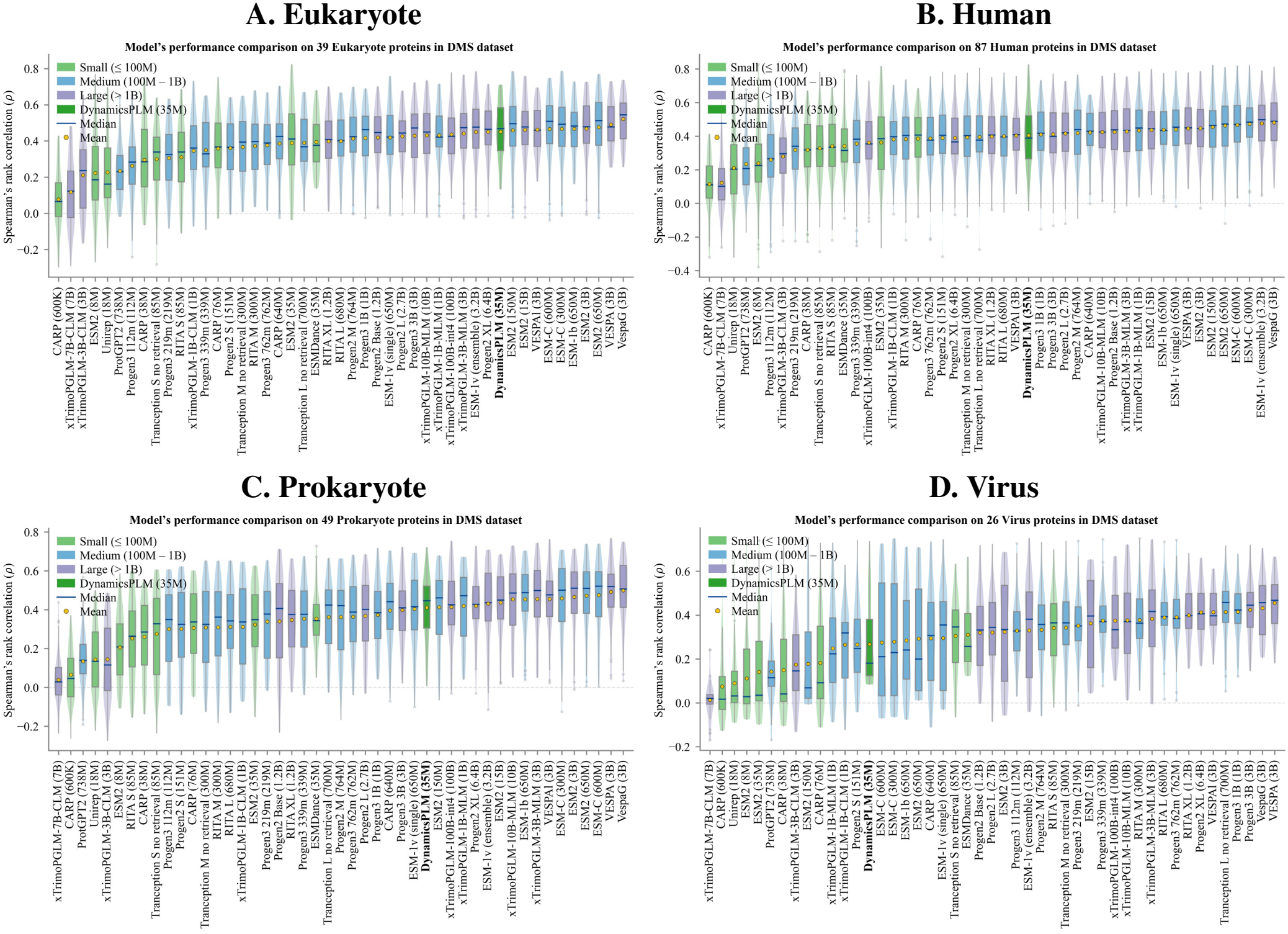
Per-taxon model performance comparison on the ProteinGym DMS substitution benchmark (47 sequence-based models, sorted by mean *ρ* within each panel). (A) Eukaryote (39 proteins). DynamicsPLM (35M): mean *ρ* = 0.451, median *ρ* = 0.469; ranks 1st among models ≤100M parameters, 11th overall. ESM2-35M (same backbone): mean *ρ* = 0.389, median *ρ* = 0.411. (B) Human (87 proteins). DynamicsPLM (35M): mean *ρ* = 0.405, median *ρ* = 0.389; ranks 1st among models ≤100M parameters, 21st overall. ESM2-35M: mean *ρ* = 0.364, median *ρ* = 0.386. (C) Prokaryote (49 proteins). DynamicsPLM (35M): mean *ρ* = 0.411, median *ρ* = 0.447; ranks 1st among models ≤100M parameters, 17th overall. ESM2-35M: mean *ρ* = 0.323, median *ρ* = 0.349. (D) Virus (26 proteins). DynamicsPLM (35M): mean *ρ* = 0.268, median *ρ* = 0.181; ranks 4th among models ≤100M parameters, 34th overall. ESM2-35M: mean *ρ* = 0.141, median *ρ* = 0.035.

## 11 PERFORMANCE COMPARISON AND ABLATION OF DYNAMICSPLM VARIANTS ON 201 PROTEINGYM DMS ASSAYS

**Table 4.**
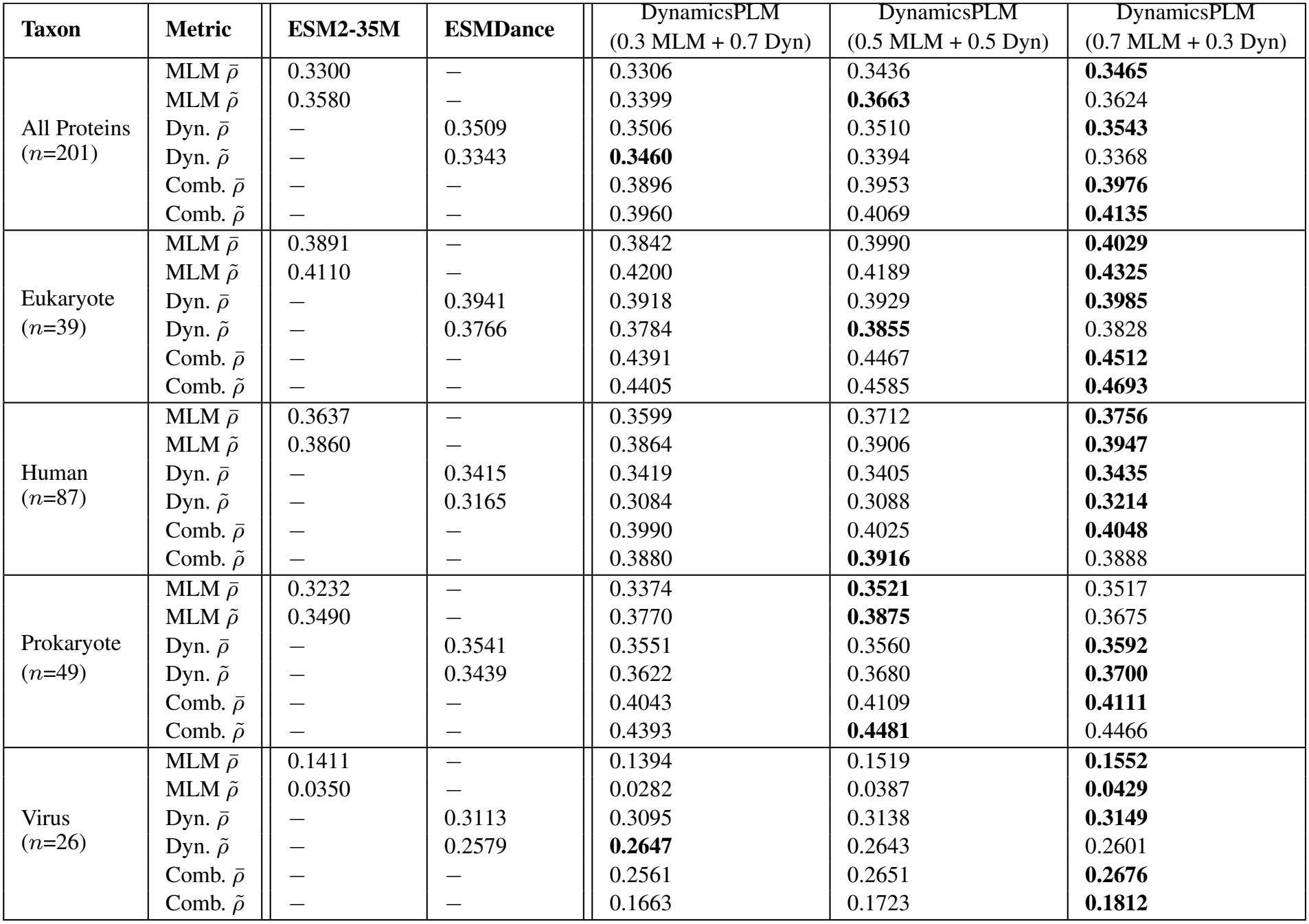
Overall performance comparison and ablation of DynamicsPLM variants on 201 ProteinGym DMS assays Notin et al. (2023), broken down by per-taxon.

| **Taxon** | **Metric** | **ESM2-35M** | **ESMDance** | DynamicsPLM<br>(0.3 MLM + 0.7 Dyn) | DynamicsPLM<br>(0.5 MLM + 0.5 Dyn) | DynamicsPLM<br>(0.7 MLM + 0.3 Dyn) |
|---|---|---|---|---|---|---|
| All Proteins<br>(*n*=201) | MLM ρ̄ | 0.3300 | — | 0.3306 | 0.3436 | **0.3465** |
|  | MLM ρ̃ | 0.3580 | — | 0.3399 | **0.3663** | 0.3624 |
|  | Dyn. ρ̄ | — | 0.3509 | 0.3506 | 0.3510 | **0.3543** |
|  | Dyn. ρ̃ | — | 0.3343 | **0.3460** | 0.3394 | 0.3368 |
|  | Comb. ρ̄ | — | — | 0.3896 | 0.3953 | **0.3976** |
|  | Comb. ρ̃ | — | — | 0.3960 | 0.4069 | **0.4135** |
| Eukaryote<br>(*n*=39) | MLM ρ̄ | 0.3891 | — | 0.3842 | 0.3990 | **0.4029** |
|  | MLM ρ̃ | 0.4110 | — | 0.4200 | 0.4189 | **0.4325** |
|  | Dyn. ρ̄ | — | 0.3941 | 0.3918 | 0.3929 | **0.3985** |
|  | Dyn. ρ̃ | — | 0.3766 | 0.3784 | **0.3855** | 0.3828 |
|  | Comb. ρ̄ | — | — | 0.4391 | 0.4467 | **0.4512** |
|  | Comb. ρ̃ | — | — | 0.4405 | 0.4585 | **0.4693** |
| Human<br>(*n*=87) | MLM ρ̄ | 0.3637 | — | 0.3599 | 0.3712 | **0.3756** |
|  | MLM ρ̃ | 0.3860 | — | 0.3864 | 0.3906 | **0.3947** |
|  | Dyn. ρ̄ | — | 0.3415 | 0.3419 | 0.3405 | **0.3435** |
|  | Dyn. ρ̃ | — | 0.3165 | 0.3084 | 0.3088 | **0.3214** |
|  | Comb. ρ̄ | — | — | 0.3990 | 0.4025 | **0.4048** |
|  | Comb. ρ̃ | — | — | 0.3880 | **0.3916** | 0.3888 |
| Prokaryote<br>(*n*=49) | MLM ρ̄ | 0.3232 | — | 0.3374 | **0.3521** | 0.3517 |
|  | MLM ρ̃ | 0.3490 | — | 0.3770 | **0.3875** | 0.3675 |
|  | Dyn. ρ̄ | — | 0.3541 | 0.3551 | 0.3560 | **0.3592** |
|  | Dyn. ρ̃ | — | 0.3439 | 0.3622 | 0.3680 | **0.3700** |
|  | Comb. ρ̄ | — | — | 0.4043 | 0.4109 | **0.4111** |
|  | Comb. ρ̃ | — | — | 0.4393 | **0.4481** | 0.4466 |
| Virus<br>(*n*=26) | MLM ρ̄ | 0.1411 | — | 0.1394 | 0.1519 | **0.1552** |
|  | MLM ρ̃ | 0.0350 | — | 0.0282 | 0.0387 | **0.0429** |
|  | Dyn. ρ̄ | — | 0.3113 | 0.3095 | 0.3138 | **0.3149** |
|  | Dyn. ρ̃ | — | 0.2579 | **0.2647** | 0.2643 | 0.2601 |
|  | Comb. ρ̄ | — | — | 0.2561 | 0.2651 | **0.2676** |
|  | Comb. ρ̃ | — | — | 0.1663 | 0.1723 | **0.1812** |

## 12 PERFORMANCE COMPARISON AND ABLATION OF DYNAMICSPLM VARIANTS ON TSUBOYAMA DATASET

**Table 5.** Overall performance comparison and ablation of DynamicsPLM variants on in Tsuboyama Dataset Tsuboyama et al. (2023) (ΔΔ*G*), broken down by natural and *de novo* proteins.

| **Taxon** | **Metric** | **ESM2-35M** | **ESMDance** | DynamicsPLM<br>(0.3 MLM + 0.7 Dyn) | DynamicsPLM<br>(0.5 MLM + 0.5 Dyn) | DynamicsPLM<br>(0.7 MLM + 0.3 Dyn) |
|---|---|---|---|---|---|---|
| All Proteins<br>(*n*=412) | MLM ρ̄ | 0.3476 | — | 0.3508 | 0.3613 | **0.3665** |
|  | MLM ρ̃ | 0.3244 | — | 0.3557 | **0.3701** | 0.3690 |
|  | Dyn. ρ̄ | — | **0.4389** | 0.4379 | 0.4368 | 0.4348 |
|  | Dyn. ρ̃ | — | **0.4567** | 0.4527 | 0.4519 | 0.4478 |
|  | Comb. ρ̄ | — | — | 0.4421 | 0.4462 | **0.4478** |
|  | Comb. ρ̃ | — | — | 0.4567 | 0.4587 | **0.4589** |
| Natural Proteins<br>(*n*=266) | MLM ρ̄ | 0.4237 | — | 0.4291 | 0.4370 | **0.4394** |
|  | MLM ρ̃ | 0.4619 | — | 0.4462 | 0.4560 | **0.4627** |
|  | Dyn. ρ̄ | — | 0.4424 | **0.4512** | 0.4461 | 0.4429 |
|  | Dyn. ρ̃ | — | 0.4555 | 0.4548 | **0.4610** | 0.4569 |
|  | Comb. ρ̄ | — | — | **0.4906** | 0.4900 | 0.4892 |
|  | Comb. ρ̃ | — | — | 0.5079 | **0.5095** | 0.5085 |
| *de novo* Proteins<br>(*n*=146) | MLM ρ̄ | 0.2090 | — | 0.2081 | 0.2233 | **0.2336** |
|  | MLM ρ̃ | 0.2142 | — | 0.2069 | 0.2078 | **0.2283** |
|  | Dyn. ρ̄ | — | **0.4325** | 0.4138 | 0.4200 | 0.4199 |
|  | Dyn. ρ̃ | — | **0.4587** | 0.4350 | 0.4377 | 0.4339 |
|  | Comb. ρ̄ | — | — | 0.3538 | 0.3666 | **0.3724** |
|  | Comb. ρ̃ | — | — | 0.3652 | 0.3768 | **0.3847** |

## Notes

### Competing Interest Statement

The authors have declared no competing interest.

