## Supplemental tables and figures for "DynamicsPLM: A Dynamics-Enriched Small Protein Language Model for Protein Variant Effect Prediction"

### Supplementary Information

#### 1 TRAIN AND EVALUATION DATASET DETAILS

**Table S1.** Number of Proteins per Data Source in the Train and Evaluation Sets. We source this data from ESMDance Hou et al. (2026).

| Group | Source | Train | Evaluation |
|---|---|---|---|
| High-resolution data | ATLAS | 1,361 | 71 |
| | GPCRmd | 107 | 5 |
| | PED | 252 | 17 |
| | mdCATH | 5,067 | 312 |
| Low-resolution data | IDRome | 26,502 | 1,360 |
| | proteinflow_pdb | 23,830 | 1,244 |
| | proteinflow_sabdab | 1,713 | 118 |
| **Total** | | **58,832** | **3,127** |

#### 2 RESIDUE-LEVEL AND PAIRWISE DYNAMIC FATURES

**Table S2.** Residue-level (50) and pairwise dynamic (13) features used to enrich DynamicsPLM, with their dimensionality and data source. MD/IDR sources (ATLAS, GPCRmd, PED, mdCATH, IDRome) supply the SASA, RMSF, secondary structure, torsion angle, and pairwise contact features; Proteinflow (pdb and SAbDab structures) supplies normal-mode-analysis-derived features only. For a given protein, features unavailable from its source carry the sentinel value ($-1$) and are excluded from the loss.

| Feature | Dim | Description | Source |
|---|---|---|---|
| *Residue features* | | | |
| sasa_mean | 1 | Mean solvent-accessible surface area | MD/IDR |
| sasa_std | 1 | SASA fluctuation (std. dev.) | |
| rmsf_nor | 1 | RMSF, normalized per protein by its max | |
| ss | 8 | 8-class secondary structure distribution | |
| chi | 12 | Side-chain $\chi_1$ torsion angle distribution (12 bins) | |
| phi | 12 | Backbone $\phi$ torsion angle distribution (12 bins) | |
| psi | 12 | Backbone $\psi$ torsion angle distribution (12 bins) | |
| nma_res1 | 1 | Residue fluctuation amplitude, normal mode 1 | Proteinflow |
| nma_res2 | 1 | Residue fluctuation amplitude, normal mode 2 | |
| nma_res3 | 1 | Residue fluctuation amplitude, normal mode 3 | |
| *Pairwise features* | | | |
| vdw | 1 | Van der Waals contact frequency | MD/IDR |
| hbbb | 1 | Backbone–backbone hydrogen bond frequency | |
| hbsb | 1 | Side chain–backbone hydrogen bond frequency | |
| hbss | 1 | Side chain–side chain hydrogen bond frequency | |
| hp | 1 | Hydrophobic contact frequency | |
| sb | 1 | Salt bridge frequency | |
| pc | 1 | $\pi$–cation contact frequency | |
| ps | 1 | $\pi$-stacking (parallel) frequency | |
| ts | 1 | $\pi$-stacking (T-shaped) frequency | |
| corr | 1 | C$\alpha$ movement correlation | |
| nma_pair1 | 1 | Pairwise movement correlation, normal mode 1 | Proteinflow |
| nma_pair2 | 1 | Pairwise movement correlation, normal mode 2 | |
| nma_pair3 | 1 | Pairwise movement correlation, normal mode 3 | |

### 3 PER-TAXON MODEL PERFORMANCE COMPARISON ON PROTEINGYM DMS SUBSTITUTION BENCHMARK

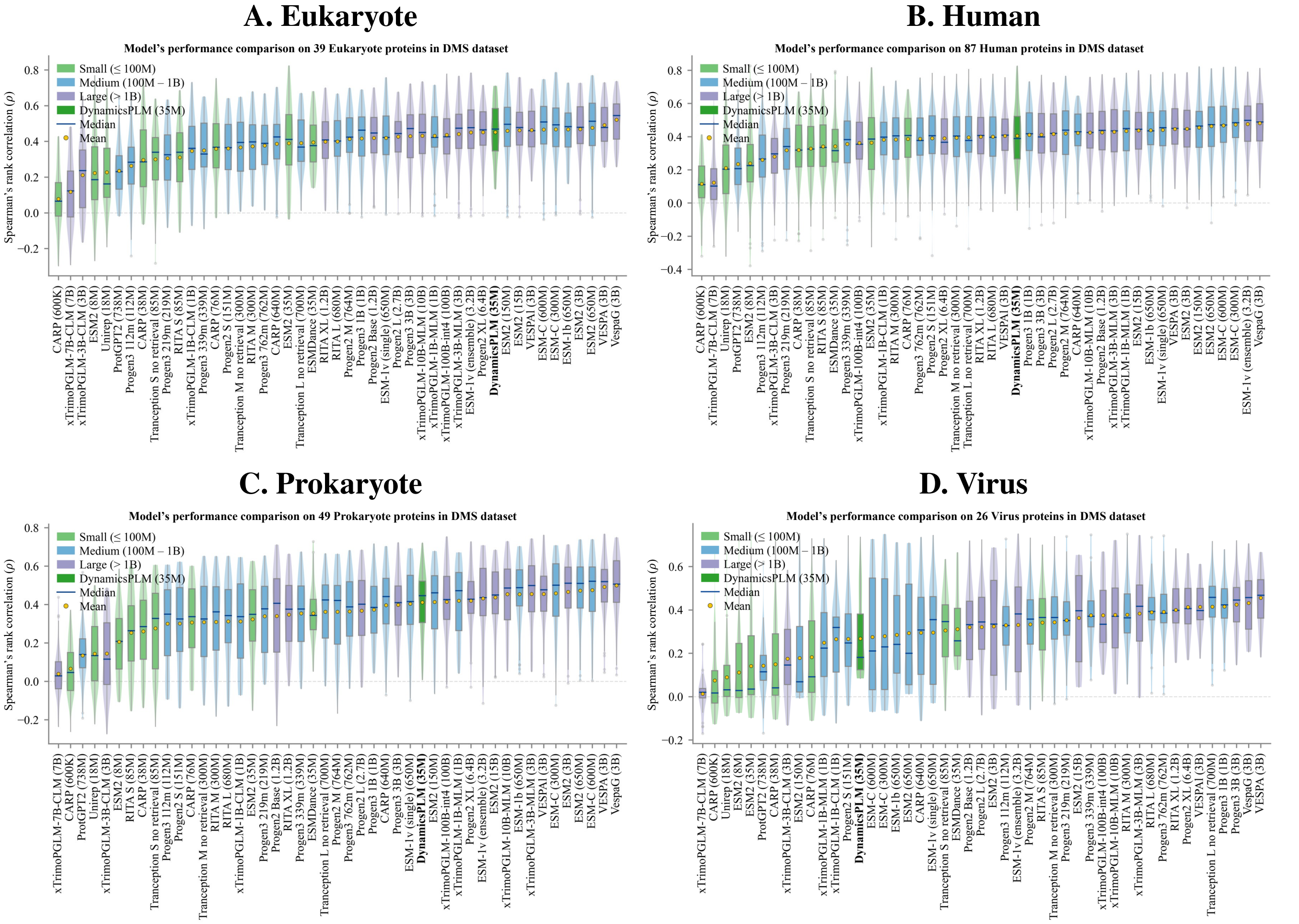


**Figure S1.** Per-taxon model performance comparison on the ProteinGym DMS substitution benchmark (47 sequence-based models, sorted by mean $\rho$ within each panel). (A) Eukaryote (39 proteins). DynamicsPLM (35M): mean $\rho = 0.451$, median $\rho = 0.469$; ranks 1st among models $\leq$100M parameters, 11th overall. ESM2-35M (same backbone): mean $\rho = 0.389$, median $\rho = 0.411$. (B) Human (87 proteins). DynamicsPLM (35M): mean $\rho = 0.405$, median $\rho = 0.389$; ranks 1st among models $\leq$100M parameters, 21st overall. ESM2-35M: mean $\rho = 0.364$, median $\rho = 0.386$. (C) Prokaryote (49 proteins). DynamicsPLM (35M): mean $\rho = 0.411$, median $\rho = 0.447$; ranks 1st among models $\leq$100M parameters, 17th overall. ESM2-35M: mean $\rho = 0.323$, median $\rho = 0.349$. (D) Virus (26 proteins). DynamicsPLM (35M): mean $\rho = 0.268$, median $\rho = 0.181$; ranks 4th among models $\leq$100M parameters, 34th overall. ESM2-35M: mean $\rho = 0.141$, median $\rho = 0.035$.

### 4 PERFORMANCE COMPARISON AND ABLATION OF DYNAMICSPLM VARIANTS ON 201 PROTEINGYM DMS ASSAYS

**Table S3.** Overall performance comparison and ablation of DynamicsPLM variants on 201 ProteinGym DMS assays Notin et al. (2023), broken down by per-taxon.

| **Taxon** | **Metric** | **ESM2-35M** | **ESMDance** | DynamicsPLM (0.3 MLM + 0.7 Dyn) | DynamicsPLM (0.5 MLM + 0.5 Dyn) | DynamicsPLM (0.7 MLM + 0.3 Dyn) |
|---|---|---|---|---|---|---|
| All Proteins ($n$=201) | MLM $\bar{\rho}$ | 0.3300 | – | 0.3306 | 0.3436 | **0.3465** |
| | MLM $\tilde{\rho}$ | 0.3580 | – | 0.3399 | **0.3663** | 0.3624 |
| | Dyn. $\bar{\rho}$ | – | 0.3509 | 0.3506 | 0.3510 | **0.3543** |
| | Dyn. $\tilde{\rho}$ | – | 0.3343 | **0.3460** | 0.3394 | 0.3368 |
| | Comb. $\bar{\rho}$ | – | – | 0.3896 | 0.3953 | **0.3976** |
| | Comb. $\tilde{\rho}$ | – | – | 0.3960 | 0.4069 | **0.4135** |
| Eukaryote ($n$=39) | MLM $\bar{\rho}$ | 0.3891 | – | 0.3842 | 0.3990 | **0.4029** |
| | MLM $\tilde{\rho}$ | 0.4110 | – | 0.4200 | 0.4189 | **0.4325** |
| | Dyn. $\bar{\rho}$ | – | 0.3941 | 0.3918 | 0.3929 | **0.3985** |
| | Dyn. $\tilde{\rho}$ | – | 0.3766 | 0.3784 | **0.3855** | 0.3828 |
| | Comb. $\bar{\rho}$ | – | – | 0.4391 | 0.4467 | **0.4512** |
| | Comb. $\tilde{\rho}$ | – | – | 0.4405 | 0.4585 | **0.4693** |
| Human ($n$=87) | MLM $\bar{\rho}$ | 0.3637 | – | 0.3599 | 0.3712 | **0.3756** |
| | MLM $\tilde{\rho}$ | 0.3860 | – | 0.3864 | 0.3906 | **0.3947** |
| | Dyn. $\bar{\rho}$ | – | 0.3415 | 0.3419 | 0.3405 | **0.3435** |
| | Dyn. $\tilde{\rho}$ | – | 0.3165 | 0.3084 | 0.3088 | **0.3214** |
| | Comb. $\bar{\rho}$ | – | – | 0.3990 | 0.4025 | **0.4048** |
| | Comb. $\tilde{\rho}$ | – | – | 0.3880 | **0.3916** | 0.3888 |
| Prokaryote ($n$=49) | MLM $\bar{\rho}$ | 0.3232 | – | 0.3374 | **0.3521** | 0.3517 |
| | MLM $\tilde{\rho}$ | 0.3490 | – | 0.3770 | **0.3875** | 0.3675 |
| | Dyn. $\bar{\rho}$ | – | 0.3541 | 0.3551 | 0.3560 | **0.3592** |
| | Dyn. $\tilde{\rho}$ | – | 0.3439 | 0.3622 | 0.3680 | **0.3700** |
| | Comb. $\bar{\rho}$ | – | – | 0.4043 | 0.4109 | **0.4111** |
| | Comb. $\tilde{\rho}$ | – | – | 0.4393 | **0.4481** | 0.4466 |
| Virus ($n$=26) | MLM $\bar{\rho}$ | 0.1411 | – | 0.1394 | 0.1519 | **0.1552** |
| | MLM $\tilde{\rho}$ | 0.0350 | – | 0.0282 | 0.0387 | **0.0429** |
| | Dyn. $\bar{\rho}$ | – | 0.3113 | 0.3095 | 0.3138 | **0.3149** |
| | Dyn. $\tilde{\rho}$ | – | 0.2579 | **0.2647** | 0.2643 | 0.2601 |
| | Comb. $\bar{\rho}$ | – | – | 0.2561 | 0.2651 | **0.2676** |
| | Comb. $\tilde{\rho}$ | – | – | 0.1663 | 0.1723 | **0.1812** |

### 5 PERFORMANCE COMPARISON AND ABLATION OF DYNAMICSPLM VARIANTS ON TSUBOYAMA DATASET

**Table S4.** Overall performance comparison and ablation of DynamicsPLM variants on in Tsuboyama Dataset Tsuboyama et al. (2023) ($\Delta\Delta G$), broken down by natural and *de novo* proteins.

| **Taxon** | **Metric** | **ESM2-35M** | **ESMDance** | DynamicsPLM (0.3 MLM + 0.7 Dyn) | DynamicsPLM (0.5 MLM + 0.5 Dyn) | DynamicsPLM (0.7 MLM + 0.3 Dyn) |
|---|---|---|---|---|---|---|
| All Proteins ($n$=412) | MLM $\bar{\rho}$ | 0.3476 | – | 0.3508 | 0.3613 | **0.3665** |
| | MLM $\tilde{\rho}$ | 0.3244 | – | 0.3557 | **0.3701** | 0.3690 |
| | Dyn. $\bar{\rho}$ | – | **0.4389** | 0.4379 | 0.4368 | 0.4348 |
| | Dyn. $\tilde{\rho}$ | – | **0.4567** | 0.4527 | 0.4519 | 0.4478 |
| | Comb. $\bar{\rho}$ | – | – | 0.4421 | 0.4462 | **0.4478** |
| | Comb. $\tilde{\rho}$ | – | – | 0.4567 | 0.4587 | **0.4589** |
| Natural Proteins ($n$=266) | MLM $\bar{\rho}$ | 0.4237 | – | 0.4291 | 0.4370 | **0.4394** |
| | MLM $\tilde{\rho}$ | 0.4619 | – | 0.4462 | 0.4560 | **0.4627** |
| | Dyn. $\bar{\rho}$ | – | 0.4424 | **0.4512** | 0.4461 | 0.4429 |
| | Dyn. $\tilde{\rho}$ | – | 0.4555 | 0.4548 | **0.4610** | 0.4569 |
| | Comb. $\bar{\rho}$ | – | – | **0.4906** | 0.4900 | 0.4892 |
| | Comb. $\tilde{\rho}$ | – | – | 0.5079 | **0.5095** | 0.5085 |
| *de novo* Proteins ($n$=146) | MLM $\bar{\rho}$ | 0.2090 | – | 0.2081 | 0.2233 | **0.2336** |
| | MLM $\tilde{\rho}$ | 0.2142 | – | 0.2069 | 0.2078 | **0.2283** |
| | Dyn. $\bar{\rho}$ | – | **0.4325** | 0.4138 | 0.4200 | 0.4199 |
| | Dyn. $\tilde{\rho}$ | – | **0.4587** | 0.4350 | 0.4377 | 0.4339 |
| | Comb. $\bar{\rho}$ | – | – | 0.3538 | 0.3666 | **0.3724** |
| | Comb. $\tilde{\rho}$ | – | – | 0.3652 | 0.3768 | **0.3847** |
